# The antiviral protein Shiftless blocks p-body formation during KSHV infection

**DOI:** 10.1101/2023.11.16.567185

**Authors:** David Hatfield, William Rodriguez, Timothy Mehrmann, Mandy Muller

## Abstract

P-bodies (PB) are non-membranous foci involved in determining mRNA fate by affecting translation and mRNA decay. In this study, we identify the anti-viral factor SHFL as a potent disassembly factor of PB. We show that PBs remain sparse in the presence of SHFL even in the context of oxidative stress, a major trigger for PB induction. Mutational approaches revealed that SHFL RNA binding activity is not required for its PB disassembly function. However, we have identified a new region of SHFL which bridges two distant domains as responsible for PB disassembly. Furthermore, we show that SHFL ability to disrupt PB formation is directly linked to its anti-viral activity during KSHV infection. While WT SHFL efficiently restricts KSHV lytic cycle, PB disruption defective mutants no longer lead to reactivation defects. SHFL-mediated PB disassembly also leads to increased expression of key anti-viral cytokines, further expanding SHFL dependent anti-viral state. Taken together, our observations suggest a role of SHFL in PB disassembly, which could have important anti-viral consequences during infection.

## INTRODUCTION

The battle between virus and host often culminates around the critical distribution of gene expression resources. A growing body of work is highlighting how viruses cleverly modulate RNA stability and localization to reshape their host cell gene expression environment. For viruses such as the gammaherpesvirus KSHV (Kaposi Sarcoma Associated Virus), during lytic replication, it is imperative to rapidly co-opt the gene expression machinery of the host and repurpose it towards viral gene expression and replication. To achieve this, KSHV induces a widespread RNA decay event that relies on its own encoded endoribonuclease SOX. This extensive RNA decay event is accompanied with numerous downstream effects including a global re-localization of RNA binding proteins (RBP) from their bound cytoplasmic mRNA targets to the nucleus^1^, a subsequent widespread shutdown of RNA pol-II based transcription^2,3^, and even hyperadenylation of nascent transcripts leading to a blockade of nuclear export ^4^. All of these events culminate into a large-scale remodeling of the gene expression environment and reallocating of resources toward viral replication while simultaneously dampening the host immune response^5,6^. Whether or how the host cell reacts and counteracts this viral takeover is not well understood.

We have previously shown that while KSHV nuclease SOX targets most of the host transcriptome, some key transcripts actively evade degradation. To date, no consensus appears to unify these escaping mRNAs with either a pro- or anti-viral function. Some, like the host Interleukin-6 (IL-6), even have well-documented pro-viral roles^7–11^. However, recently, we identified a novel escapee -SHFL-which stringently resists SOX cleavage and encodes a potent anti-viral protein that restricts KSHV Lytic replication^12,13^. SHFL is an interferon stimulated gene (ISG) that is emerging as a critical piece of the innate immune response to viral infection, capable of suppressing the replication of multiple DNA, RNA, and retroviruses^14^. Recent reports over the past 5 years revealed that SHFL can negatively modulate viral RNA stability, viral gene translation, and even viral protein stability^13,15–19^. In the context to KSHV, we showed that SHFL broadly restricts KSHV lytic gene expression, including that of the master latent-to-lytic switch protein, KSHV ORF50 (RTA), resulting in a global downregulation of the KSHV lytic gene cascade. Surprisingly, we also found that SHFL expression appears to disrupt Processing Bodies (P-bodies or PB) both outside of and in the context of KSHV infection. This capacity of SHFL to modulate cytoplasmic RNA granules represents one of the first examples of an ISG capable of influencing P-body dynamics during viral infection.

In mammalian cells, formation of RNA granules allows flexibility for survival during adverse biological conditions^20^. PB are non-membranous cytoplasmic ribonucleoprotein (RNP) foci long believed to be solely sites of RNA decay, but are now regarded as sites of RNA stasis, acting as a “storage facility” for constitutive turnover and regulated release of key transcripts into the translation pool^21–26^. While PBs are constitutively present in most cell types, they are also dynamic, changing in size and number in response to shifts in cellular environmental conditions and gene expression^27–31^. This plasticity of PBs as biomolecular condensates has been attributed to their ability to undergo liquid-liquid phase separation. PB formation occurs via the sequential accumulation of multivalent RNP complexes, often enriched with RBP constituents that contain intrinsically disordered regions that serve as essential scaffolds for PB maturation. Excitingly, recent lines of research have begun to uncover cellular factors that drive the regulation of PB assembly and fission. While numerous viral proteins have been shown to modulate PB dynamics, to-date, there are very few examples of cellular proteins that actively drive the disassembly of PB in human cells^32,33^.

Several studies have shown that the RNA composition of P-bodies often includes proinflammatory cytokine transcripts such as Interleukin-6 (IL-6), Tumor Necrosis Factor (TNF), and Interleukin-8 (CXCL8)^29,34^. The finely regulated timing of the translation and turnover of these transcripts - especially when the cell is not under threat by pathogens - is critical to maintaining cellular homeostasis^34,35^. This is exemplified by the fact that many of these same mRNA bear RNA elements located in their 3’ UTR, known as AU-rich elements (ARE), that can actively signal their sequestration into PBs^36–38^. Thus, the loss of PBs has been hypothesized to be an antiviral strategy, priming a robust defense for the cell^39^. However, to date, there are scarce examples of how the cell might induce P-body loss to facilitate this process^12,32,33^. Complicating this model further are an increasing number of reports that viral infection by several DNA and RNA viruses induce P-body loss via viral gene products that directly interfere with the scaffolding of PB-associated RNPs^36,39–42^.

Given its significant role in the innate immune response, we hypothesized that SHFL impact on PB dynamics could serve as a host driven defense mechanism during KSHV infection. Here, we show that SHFL directly blocks PB formation, even in the context of chemically-induced stress. We also found that SHFL binding RNA is not required in its induction of PB loss. Instead, we define amino acids 151 to 200 and 190 to 241 as a new domain of SHFL critical for this disruption. As expected, SHFL-induced PB disruption also led to an increase in inflammatory cytokine expression which could further bolsters SHFL antiviral capacity. Reinforcing this idea, we show that SHFL ability to disrupt PB is linked to its anti-KSHV activity as the ΔPB mutant is no longer able to restrict KSHV lytic replication. Collectively our results demonstrate that SHFL acts as a regulator of PB dynamics during herpesvirus infection, restricting their formation to promote the expression of proinflammatory cytokines and contribute to a global antiviral state.

## RESULTS

### SHLF blocks PB formation

We recently showed that during KSHV infection, upon expression of the anti-viral factor SHFL, cytoplasmic RNA granules labeled as P-bodies (PB) are lost^12^. Moreover, we now observe that SHFL expression alone, even outside of the context of infection, leads to PB loss [**FIG1A**], therefore ruling out the hypothesis that this may be triggered by a KSHV protein. Given the stringent reduction in PB numbers in cells overexpressing SHFL, we next sought to understand if SHFL blocks *de novo* formation of PB or triggers their dissolution. To test this, we treated cells with sodium arsenite (NaAs), a known oxidative stressor for cells that induces a marked PB increase [**FIG 1B**]^43^. A gradient of NaAs treatment was performed to best identify the concentration that would significantly increase PB counts yet minimized cytotoxicity and abnormalities in cell morphology [**FIG S1**]. Treatment was performed over a period of 30 min before fixing the cells and staining with the PB marker DDX6. Upon treatment with 0.25mM NaAs, we observed an increase in PB numbers as expected. This treatment was then performed in cells expressing SHFL: if SHFL blocks *de novo* formation of PB, then we would expect to see no increase PB numbers; however, if SHFL triggers the disassembly of PB, we would expect to still see a large increase of PB after 30 min. In line with our hypothesis, we observed that when SHFL is overexpressed, NaAs-treated cells no longer display higher PB counts, suggesting that SHFL is actively prevents their formation.

**Figure 1:**
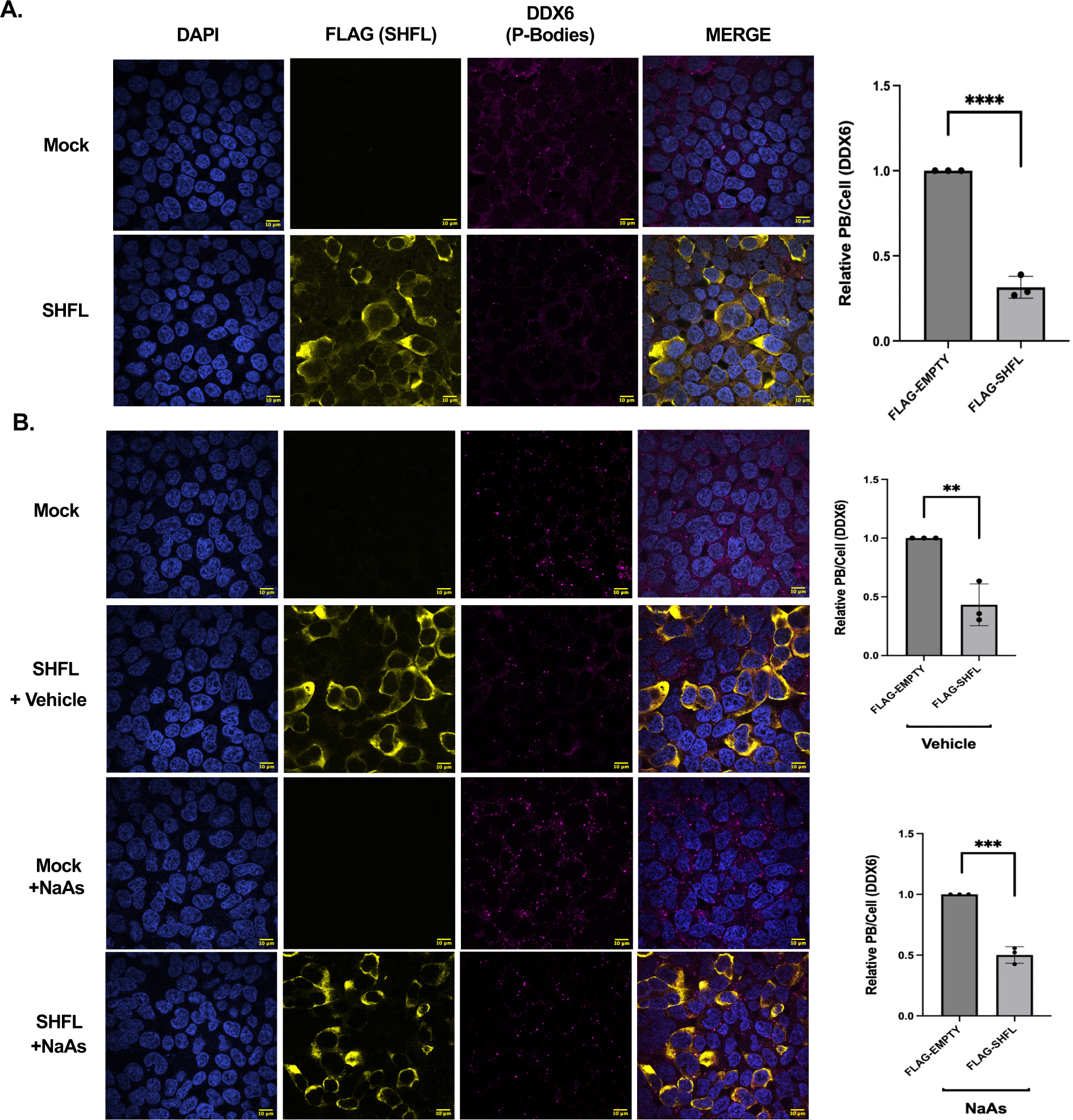
SHFL blocks de novo synthesis of PB. **(A)** HEK293T cells were transfected with either a Flag-tagged SHFL (or mock). 24h later, cells were subjected to immunofluorescence assay and stained for the indicated processing body marker (DDX6). **(B)** HEK293T cells were transfected with either a Flag-tagged SHFL (or mock). 24h later, cells were treated for 30 min with 25mM NaAs then subjected to immunofluorescence assay and stained for the indicated processing body marker (DDX6). The number of P-body puncta per cell was quantified using the CellProfiler pipeline as described in the methods and normalized to the mock control within each replicate. Here and in all figures, scale bar represents 10 μm and statistics were determined using students paired t-test between control and experimental groups; error bars represent standard error of the mean; n=3 independent biological replicates. n.s.=not significant, **, P<0.01, ***, P < 0.001.

### SHFL-mediated PB loss does not rely on SHFL RNA binding activity

PB bodies are condensates composed of RNA and proteins. SHFL has previously been shown to bind to mRNA so we hypothesized that SHFL could impede PB formation by binding and sequestering key RNA molecules away from PB, preventing their assembly. To test this hypothesis, we constructed mutants to abrogate SHFL RNA binding activity: one mutant that lacks amino acids 102-150 (ΔRBD), hypothesized to contain the SHFL RNA-binding domain and previously reported to be unable to bind to mRNA^44^. We also generated a second SHFL mutant introducing point mutations at arginine 131, 133 and 136 (Δ3R) that have been implicated as the critical amino acids responsible for SHFL RNA binding *in-vitro*. We first verified that these mutants expressed properly [**FIG2A**] and then proceeded to quantify PB content in cells expressing each of them. We observed that similar to WT SHFL, these mutants also drastically reduced PB count, suggesting that SHFL RNA binding activity is not required for PB disruption.

**Figure 2:**
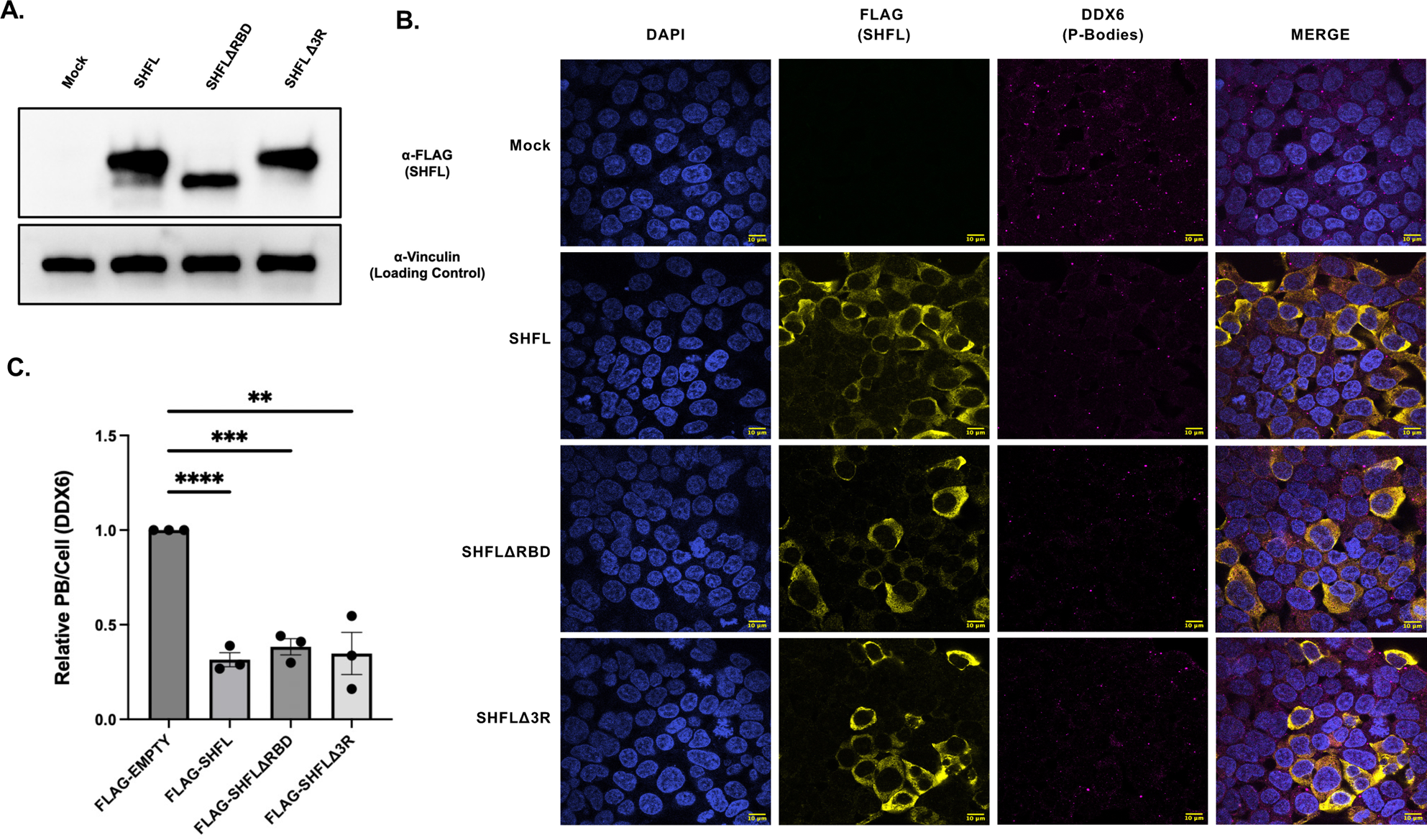
SHFL RNA binding activity is not required for PB disruption. **(A)** HEK293T cells were transfected with either a full-length Flag-tagged SHFL or SHFL RNA binding domain mutants (ΔRBD or Δ3R as indicated). 24h later, cells were harvested and subjected to immunoblot and stained with the indicated antibodies. **(B-C)** HEK293T cells were transfected with the indicated SHFL constructs. 24h later, cells were subjected to immunofluorescence assay and stained for the indicated processing body marker (DDX6). The number of P-body puncta per cell was quantified using the CellProfiler pipeline as described in the methods and normalized to the mock control within each replicate.

### SHFL bridge domain is responsible for PB loss

We next sought to identify the domain in SHFL that may trigger PB loss and thus constructed a library of SHFL mutants with incremental 50 amino acid deletions from the N-terminus. As shown in **figure 3**, two domain deletion mutants appear to result in the restoration of PB numbers, suggesting that individually, these domains contribute to the disruption of PB. We therefore focused on these two domains and constructed deletion mutants for each domain (Δ151-200 and Δ251-291) individually or in combination. However, deleting both domains at once did not result in an additive effect on PB restoration (**figure 4**). Thus, we hypothesized that these domains could broadly affect SHFL subcellular localization. Yet both single deletion mutant and double deletion mutant have a localization like that of WT SHFL and are present in the cytoplasm (**fig 4B**). Of note, deleting SHFL NES (Nuclear Export Signal) which is predicted to be localized within the C-terminus of SHFL did not affect SHFL ability to restrict PBs. Given that these two domains are not contiguous, it was puzzling to understand how and why they could both impact SHFL-mediated PB restriction. To better understand this, we generated a structural model of WT SHFL in Alpha Fold2 (**Fig 4D**), and we were surprised to see that these domains are in-fact predicted to be in close proximity creating a common interface that we hypothesized could come together as a binding platform. In fact, we noticed two amino acids (G259 and W191) that appear to make contact between these two domains. We thus next mutated these amino acids into alanine to abrogate bond formation (referred to as ΔGW). Using this mutant, we demonstrate that SHFL ability to restrict PB formation in-fact stems from this bridge region formed by these two residues, further suggesting that SHFL may need to bind to a specific interactor to prevent PB formation.

**Figure 3:**
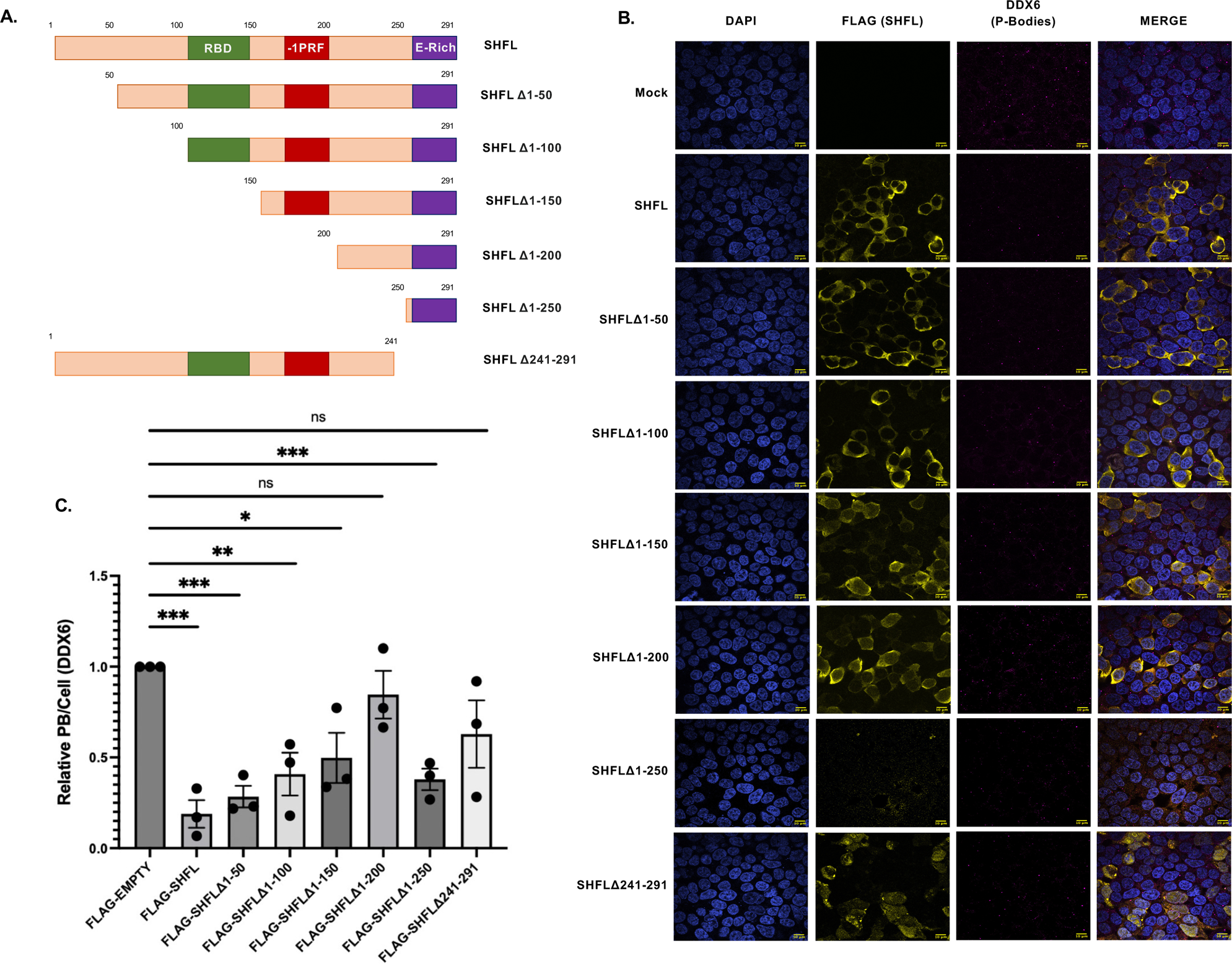
Several domains of SHFL contribute to PB restriction. **(A)** schematic representation of SHFL incremental deletion mutants. **(B)** HEK293T cells were transfected with the indicated Flag-tagged SHFL constructs and 24h later, cells were harvested and subjected to immunoblot and stained with the indicated antibodies. (**C-D**) Cells were transfected with the indicated SHFL constructs. 24h later, cells were subjected to immunofluorescence assay and stained for the indicated processing body marker (DDX6). The number of P-body puncta per cell was quantified using the CellProfiler pipeline as described in the methods and normalized to the mock control within each replicate.

**Figure 4:**
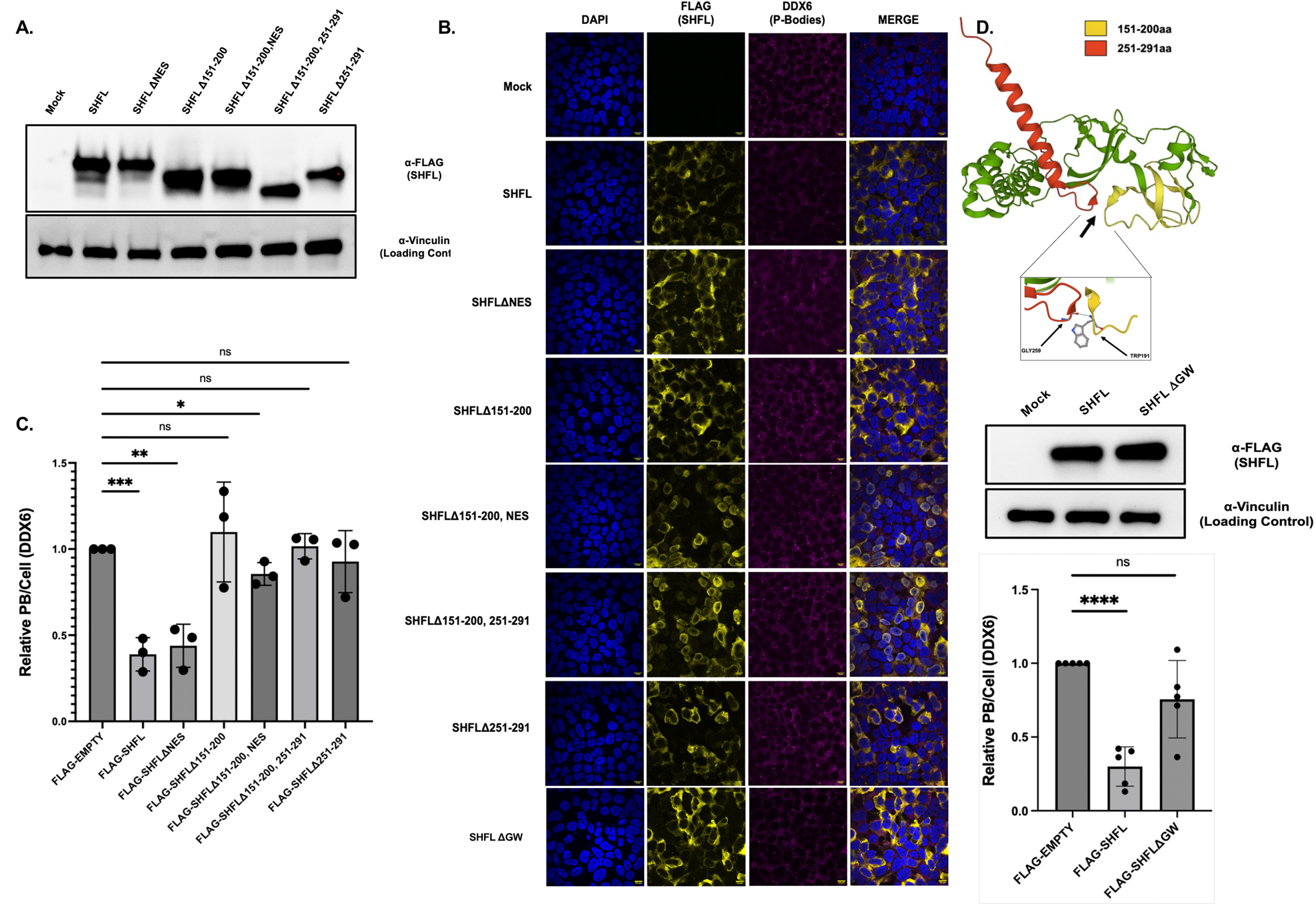
Domains 151-200 and 241-290 form an interface important necessary for PB restriction. (**A**) HEK293T cells were transfected with the indicated Flag-tagged SHFL constructs and 24h later, cells were harvested and subjected to immunoblot and stained with the indicated antibodies. (**B-C**) Cells were transfected with the indicated SHFL constructs. 24h later, cells were subjected to immunofluorescence assay and stained for the indicated processing body marker (DDX6). The number of P-body puncta per cell was quantified using the CellProfiler pipeline as described in the methods and normalized to the mock control within each replicate. (**D**) Alpha fold prediction of SHFL folding highlighting closeness of domain 151-200 and 241-290, and amino acid G259 and W191 possible bond. G259A-W191A (ΔGW) mutant of SHFL was constructed and transfected into HEK-293T cells, confirming that these mutations did not affect the stability SHFL. Immunofluorescence of stained PB however reveal no loss of PB upon expression of this mutant.

### SHFL-mediated PB loss is required to restrict KSHV lytic replication

SHFL acts as a potent anti-viral factor during KSHV infection, and we hypothesized that its role on PB repression may further contribute to establishing an anti-viral state in cells. PB are often implicated as sites of translational arrest and RNA storage. The loss of PB upon SHFL expression could thus alter the availability of certain transcripts. AU-rich elements (ARE) containing transcripts, often encode for pro-inflammatory cytokines and can thus be tied to the inflammatory response and have been found to be modulated by PB abundance. We wanted to test if SHFL expression and subsequent PB loss would also result in alteration of ARE mRNA levels. To test this we next analyzed the abundance of select ARE-mRNAs and found that several of these genes had increased transcript levels upon SHFL overexpression in uninfected cells (**Figure 5**). Given this global anti-viral potential due to SHFL-mediated PB restriction, we next wondered if SHFL capacity to restrict the KSHV lytic cycle could be attributed to its ability to inhibit PB formation. To this end, we first wanted to confirm the contribution of endogenous SHFL expression to PB numbers during KSHV infection. Upon knockdown of SHFL in iSLK.219 cells, as previously observed, there is a marked increase in KSHV lytic replication as indicated by increased RFP fluorescence^13^. This corresponded with a marked increase in P-body numbers observed in both KSHV latent and lytic cells. Next, we measured the global effect of SHFL on KSHV lytic reactivation using complementation approach the KSHV.219 model. As expected, after knocking down endogenous SHFL levels, overexpression of WT SHFL led to a drastic reduction in RFP positive cells upon KSHV reactivation, confirming SHFL anti-viral role during infection. However, SHFL mutant Δ251-291, and ΔGW which is unable to restrict PB formation also did not restrict KSHV lytic cycle. Our results therefore suggest that SHFL anti-KSHV activity at least partially results from its ability to modulate PB formation and thus to globally reshape the infected cells environment.

**Figure 5:**
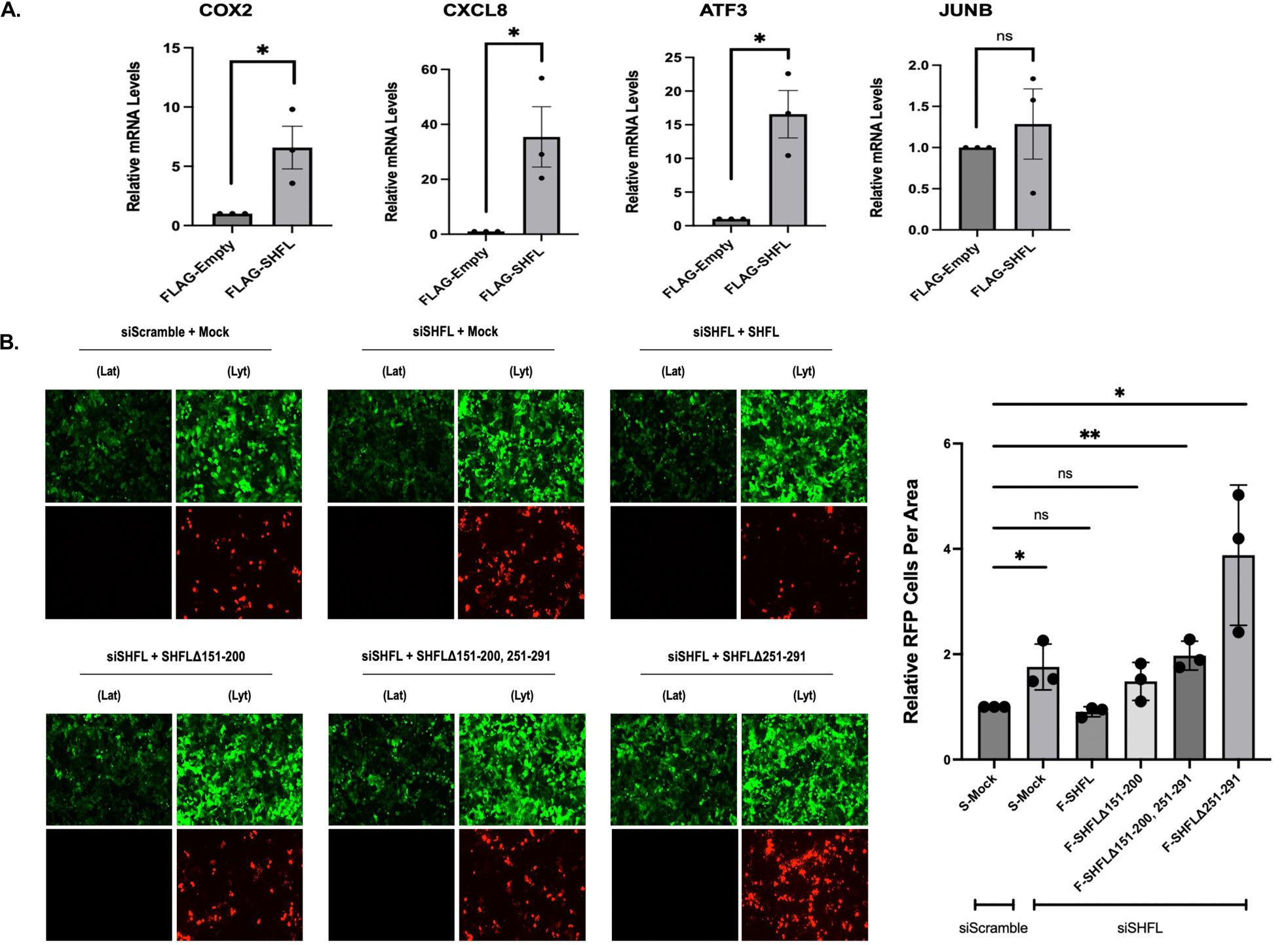
PB loss upon SHFL expression results in ARE mRNA overexpression and KSHV lytic restriction. **(A)** Cells were transfected with SHFL WT or Δ251-290. Total RNA was then harvested and subjected to RT-qPCR to measure mRNA levels of the indicated ARE-containing mRNA. (**B**) KSHV positive iSLK.219 cells were either treated with siRNAs targeting SHFL (or control non-target siRNAs - siSCRAMBLE) for 48h then transfected with the indicated Flag-tagged SHFL. Cells were then reactivated with doxycycline and sodium butyrate for 24h, then checked for reactivation efficiency by monitoring the expression of GFP and RFP. Quantification of RFP-positive cells is to the right. Values represent at least five independent views of the infected cells.

## DISCUSSION

The ability of the cell to regulate the fate of RNA in response to viral infection is crucial for their survival and adaptation. As such, both cell and virus have evolved a myriad of RNA binding proteins (RBPs) that clash for control of post-transcriptional surveillance of RNA trafficking, translation, and stability. In this context, RNA granules have emerged discreet sites of intense RNA regulation, and thus of particular hijacking during infection. The condensation of RNA into P-bodies is especially impactful, as many of the RNA species that constitute this granule type have been tied to critical cellular processes such as the pro-inflammatory response. Indeed, several previous studies have shown an enrichment of ARE-bearing cytokine mRNA within P-bodies of several cell types. These observations imply that that cell may utilize P-body formation as a means of preventing the premature translation of these transcripts or to careful time their turnover in an ever-changing cellular environment. This also suggests that properly coordinated disassembly of P-bodies may correlate to a coordinated inflammatory gene expression cascade that would ultimately promote cell survival and signaling to the greater cell microenvironment. It thus perhaps unsurprising that several viruses have evolved to block the formation of, or degrade, P-bodies. In this study, we focus on the gammaherpesvirus KSHV, a virus that has been shown to restrict PBs early-on following lytic reactivation from latency ^36,42,45,46^. It should be noted that this restriction was also observed in KSHV latent cells, but this phenotype appears to vary between latent cell types. During lytic replication, this restriction is mediated by the early lytic gene ORF57, which blocks de-novo P-body assembly by interfering with interaction between critical P-body scaffolding factors such as GW182 and Ago2 ^42^. This suggests KSHV and other viruses may actively disassemble PB to facilitate replication at the expense of triggering the host cell cytokine response. However, as is well understood for KSHV, this could contribute to establish a pro-tumorigenic environment. Other viruses appear to also benefit from PB disruption, as was shown recently by Kleer whereby dispersed PBs during Coronavirus infection lead to an upregulation of cytokine mRNA stability and translation^39^. The authors propose that this may be a mechanism that supports a pro-viral dysregulation of the cytokine response and therein ultimately promotes coronavirus pathogenesis. Collectively, these reports highlight the need for a better understanding of how the host cell differentially regulates these RNP condensates to control gene expression dynamics during viral infection.

Here, we focus on the host protein SHFL, that we previously identified as a transcript that actively evade herpesvirus-induced RNA decay^12–14^. This broad escape from viral endonuclease cleavage mirrors SHFL own broad antiviral capacity, as SHFL can restrict the replication of multiple DNA, RNA, and retroviruses through diverse mechanisms ^14^. Upon further investigation we found that following this escape from cleavage, the SHFL protein goes on to impede KSHV lytic reactivation from latency and broadly suppress viral gene expression. This potent antiviral capacity appears to stem from a direct restriction of the expression of the master regulator of KSHV RNA fate, KSHV ORF57^12^. Intriguingly, we also found that SHFL expression restricts PB numbers both outside of and within the context of KSHV infection. Little is known about cellular factors that have the capacity to disassemble PB^32,33^. As such, here, we set out to investigate the molecular underpinnings of the influence of SHFL over PBs, hypothesizing that it may be tied to the expression of pro-inflammatory cytokines and KSHV lytic replication. To our surprised, we found that SHFL restricts PBs even under Sodium Arsenite stress. Given the timing of this restriction, we concluded that SHFL must be blocking their aggregation as opposed to triggering their disassembly. Given that SHFL is a known RNA-binding protein, we thought that the ability to SHFL to bind to RNA was required for this capacity to impede PB assembly, perhaps by blocking the incorporation of specific RNA species into these phase-separated condensates. To test this, we generated mutants of SHFL that have been previously shown to oblate SHFL RNA interactions but we found that the ability of SHFL to bind to RNA is not required for its ability to restrict PB formation. A recent report of identified another host protein, Sbp1, was shown to trigger PB disassembly relied on a direct protein-protein interaction with a decapping enzyme. This suggested that SHFL roles in blocking PB formation can perhaps also stem from its protein network. We have previously mapped SHFL interaction network, and found several E3 ligases^32^. It would be interesting to test if these interactions are needed for SHFL-induced PB restriction.

To better understand which domain of SHFL was required for restriction of PB formation, we generated a SHFL deletion mutant library and tested their capacity to restrict P-body numbers. From this screen, we identified two domains that individually contribute to the disruption of PB including aa151-200 and the C-terminal end of SHFL aa251-291. Previous work by Suzuki *et al.* and others has demonstrated that there is a functional Nuclear Export Signal (NES) within the SHFL C-terminus^16^. Given the nearly exclusive cytoplasmic localization of SHFL across multiple cell types, we thought that perhaps the loss of these domains may obstruct proper SHFL subcellular localization. However both the single domain and double domain deletion mutants still demonstrate cytoplasmic localization like wild-type (WT) SHFL. This was a puzzling result, as these two domains of SHFL are not contiguous. Thus, we turned to predictive modeling of SHFL structure *via* application of Alphafold 2 to gain further structural insights in the absence of a current SHFL crystal structure. To our surprise, we found that these two regions come into close proximity in the 3D structure and there is an amino acid pair (G259 and W191) that bridges these distinct SHFL domains. Mutagenesis of these residues resulted in a SHFL unable to restrict PB formation, confirming that this is in fact the stabilization of these two domains that is necessary for PB restriction.

Lastly, we wanted to determine the contribution of SHFL-mediated suppression of PB over KSHV lytic replication. PB have been known to house pro-inflammatory mRNA transcripts that contain AU-rich (ARE) elements including Interleukin-6 (IL-6) and CXCL8 ^29,34^. The release of these transcripts from PB has been tied to a direct enhancement of mRNA stability and translation. Concordantly, we found that SHFL expression does in-fact lead to increased mRNA levels of multiple pro-inflammatory cytokines both outside the context of viral infection and during KSHV latency and lytic replication. This effect on these specific cytokine mRNA levels is likely post-transcriptional, as we did not detect any transcriptional increase by 4SU labelling of nascent RNA for these cytokines. This increase in global cytokine production would undoubtedly contribute to a global anti-viral state instated by SHFL expression. Thus, we next wanted to confirm that the influence of SHFL over PB contributes to its capacity to restrict KSHV lytic replication following reactivation from latency. Accordingly with our previous findings, we found that loss of both functional domains aa151-200 and the SHFL C-terminus aa251-291 led to a marked defect in restricting KSHV lytic replication when compared to WT SHFL. All together, these results suggest that the ability of SHFL to restrict PB directly contributes to its ability at least in part, to restrict KSHV lytic replication.

One critical factor of our investigation of SHFL and KSHV is the timing of SHFL expression. In our pilot study of SHFL, we found that the expression of SHFL climbs over the course lytic replication^13^. This makes sense from the host perspective, as SHFL mRNA has evolved to escape KSHV-mediated RNA decay and therein contribute to restrict herpesviral replication. Therefore, our observation that SHFL expression climbs over the course of KSHV lytic replication may represent a gradual suppression of PB by the host to combat KSHV infection. However, alternatively, the expression of SHFL may be considered a mechanism promoted by KSHV to promote PB disassembly in synchrony with virus encoded factors such as ORF57 and KapB. Thus, it appears that control of the timing of PB disassembly and how host factors such as SHFL or viral factors such as ORF57 and KapB may come together to dictate the ultimate outcomes of the inflammatory response to KSHV infection. For an oncogenic virus like KSHV, knowing more about the long-term consequences of this viral-host battle for the control of PB will be critical to better decipher pathways involved in both intracellular and paracrine signaling during tumorigenesis.

In summary, we identify SHFL as among the few cellular genes capable of inducing P-body disassembly and that this influence over RNA granule dynamics contributes, at least in part, to its capacity to restrict KSHV infection. It remains to be understood what host factors SHFL needs to interact with to facilitate the molecular underpinnings. Our results highlight a tight interplay between host factors like SHFL and viral factors like ORF57 for controlling PB during KSHV infection. This relationship will undoubtedly shed light on how SHFL establishes a global antiviral state. The work presented here highlights the efforts by the host cell to not only combat viral infection, but also actively modulate the fate of specific mRNA through the shield of innate immune signaling. Furthermore, our findings underline a need to better understand how the host cell regulates RNA granule disassembly and formation to mediate a versatile control over broader cellular response to a diverse range of environmental challenges and oncogenesis.

## MATERIALS AND METHODS

### Cells and transfections

HEK293T cells (ATCC) were grown in Dulbecco’s modified Eagle’s medium (DMEM; Invitrogen) supplemented with 10% fetal bovine serum (FBS). The KSHV-infected renal carcinoma human cell line iSLK.BAC16 (iSLK.WT) (kind gift from Dr. B. Glaunsinger) bearing doxycycline-inducible RTA was grown in DMEM supplemented with 10% FBS. KSHV Lytic reactivation was induced by the addition of 1 μg/ml doxycycline (BD Biosciences) and 1 mM sodium butyrate for 48 hr as reported above. For DNA transfections, cells were plated and transfected after 24 h when 70% confluent using PolyJet (SignaGen). For small interfering RNA (siRNA) transfections, cells were reverse transfected in 6-well plates by INTERFERin (Polyplus Transfection) with 10 μM siRNAs. siRNAs were obtained from IDT as Dicer-substrate siRNA (DsiRNA; siRNA C19ORF66, hs.Ri.C19orf66.13.1). NaAs treatment was performed on cells were grown on coverslips, transfected as above, and treated with NaAs (0.25mM) for 30min incubated at 37°C. Following treatment, cells were harvested and fixed via 4% Paraformaldehyde.

### Plasmids

The C19ORF66 coding region was obtained as a gBlock from IDT and cloned in a pcDNA4 Nter-3×FLAG vector (FLAG-SHFL). A FLAG-mock (pcDNA4 Nter 3xFLAG) was used as a control where indicated. The SHFL mutant library was cloned by truncating 50aa starting from the N-terminus. From the full-length SHFL, the FLAG-SHFLΔRBD mutant was constructed to lack amino acids 102-150 (FLAG-SHFLΔ102-150). The FLAG-SHFL Δ3R was ordered as a G-Block from IDT with the 3 point mutations R131A, R133A, and R136A and cloned into a pcDNA4 Nter-3xFlag vector. The mutants FLAG-SHFLΔNES and FLAG-SHFLΔ151-200 + NES were also ordered from IDT as G-Blocks with the point mutations L261A, L264A, L267A, and L269A and cloned into pcDNA4 Nter-3xFlag vectors. FLAG-SHFLΔ151-200, FLAG-SHFLΔ151-200, 251-291, FLAG-SHFLΔ251-291, and FLAG-SHFLΔW191A, G259A were ordered as G-Blocks from IDT and cloned into a pcDNA4 Nter-3xFlag vector. G-Blocks were cloned into the vector backbones via in-fusion cloning (Clonetech-takara); all final constructs were verified via Sanger Sequencing.

### RT-PCR

Total RNA was harvested using TRIzol according to the manufacture’s protocol. cDNAs were synthesized from 1 μg of total RNA using AMV reverse transcriptase (Promega) and used directly for quantitative PCR (qPCR) analysis with the SYBR green qPCR kit (Bio-Rad). Signals obtained by qPCR were normalized to 18S unless otherwise noted. qPCR Primers used in the study are as follows: COX2 (Fwd: CCCTTGGGTGTCAAAGGTAA; Rv: GCCCTCGCTTATGATCTGTC); CXCL8(Fwd: AAATCTGGCAACCCTAGTCTG; Rv: GTGAGGTAAGATGGTGGCTAAT); ATF3 (Fwd: CAAGAACGAGAAGCAGCATTTG; Rv: TTCTGGAGTCCTCCCATTCT); JUNB (Fwd: TCTACCACGACGACTCATACA; Rv: GGCTCGGTTTCAGGAGTTT).

### Immunoblotting

Cell lysates were prepared in lysis buffer (NaCl, 150 mM; Tris, 50 mM; NP-40, 0.5%; dithiothreitol [DTT], 1 mM; and protease inhibitor tablets) and quantified by Bradford assay. Equivalent amounts of each sample were resolved by SDS-PAGE and immunoblotted with each respective antibody in TBST (Tris-buffered saline, 0.1% Tween 20). Antibodies used are Mouse anti-Flag (Invitrogen, 1:1000); Rabbit anti-DDX6 (Invitrogen, 1:100) or Mouse anti-DDX6 (Sigma, 1:100); and Rabbit anti-Vinculin (Invitrogen 1:1000). Primary antibody incubations were followed by horseradish peroxidase (HRP)-conjugated goat anti-mouse or goat anti-rabbit secondary antibodies (1:5,000; Southern Biotechnology).

### Immunofluorescence

HEK293T or iSLK.WT cells were grown on coverslips and fixed in 4% formaldehyde for 20 min at room temperature. Cells were then permeabilized in 1% Triton X-100 and 0.1% sodium citrate in phosphate-buffered saline (PBS) for 10 min, saturated in bovine serum albumin (BSA) for 30 min, and incubated with the designated antibodies. After 1 h, coverslips were washed in PBS and incubated with Alexa Fluor 680, 594 or 488 secondary antibodies at 1:1,500 (Invitrogen). Coverslips were washed again in PBS and mounted in DAPI-containing Vectashield mounting medium (Vector Labs) to stain cell nuclei before visualization by confocal microscopy on a Nikon A1 resonant scanning confocal microscope (A1R-SIMe). The microscopy data were gathered in the Light Microscopy Facility and Nikon Center of Excellence at the Institute for Applied Life Sciences, UMass Amherst, with support from the Massachusetts Life Sciences Center.

### RNA Granule Quantification

Processing bodies were quantified using an unbiased image analysis pipeline generated in the freeware CellProfiler (cellprofiler.org). Briefly, detection of nuclei in the DAPI channel image was performed by applying a binary threshold and executing primary object detection between 40 and 200 pixels. From each identified nuclear object, the “Propagation” function analyzes the 488nm channel image (identifying FLAG-SHFL) to identify transfected cell borders. The pipeline then subtracts out the nuclei from the total area to define the cytoplasmic spaces. Using this cytoplasmic mask, the 594nm channel image, with the fluorescing P-body puncta (stained with DDX6), could be overlayed to only identify cytoplasmic foci (i.e. excluding any nuclear signals). The “Enhance Speckles” function reduced background staining in the cytoplasmic channel. The “global thresholding with robust background adjustments” function specifically defines and denotes certain puncta based upon strict size and intensity ranges, to reduce the amount of background artifacts quantified. All size and intensity ranges and thresholds throughout the pipeline remained unchanged between experiments. Those puncta of specific size and intensity were quantified and used for data analysis.

### Statistical analysis

All results are expressed as means ± standard errors of the means (SEMs) of experiments independently repeated at least three times. Unpaired Student’s *t* test was used to evaluate the statistical difference between samples. Significance was evaluated with *P* values as follows: * p<0.05; ** p<0.01; *** p<0.001.

## Acknowledgments

We thank all members of the Muller lab for helpful discussions and suggestions. Special thanks to Dr. Britt Glaunsinger for both our iSLK cell lines and KSHV antibodies. We are also extremely grateful to Dr. Jennifer Corcoran for her help regarding PB quantification and biology. We would also like to thank Dr. James Chambers at the UMASS IALS Light Microscope Facility for their help with protocol development and data acquisition.

